# Revisiting Tumor Mutational Burden Cutoff: Multi-Study Replicability in Immunotherapy

**DOI:** 10.1101/2025.01.02.631104

**Authors:** Iman Jaljuli, Karissa Whiting, Evan Rosenbaum, Li-Xuan Qin

## Abstract

**PURPOSE:** Tumor Mutational Burden (TMB) is a crucial biomarker for predicting the effectiveness of cancer immunotherapy. However, ongoing debates about the optimal cutoff significantly impact the clinical application of immunotherapy. The purpose of our study is to comprehensively evaluate TMB cutoffs for predicting immunotherapy outcomes across multiple studies, considering both cancer type and outcome endpoint, using statistically principled approaches.

**METHODS:** We analyzed data from PredictIO, curated through a PubMed search for studies involving immune checkpoint blockade treatment in the adjuvant setting, excluding those involving combinations with chemotherapy, targeted treatment, or radiation. The included studies, published between January 2015 and June 2022, required tumor sequencing data from whole exomes or targeted gene panels for TMB assessment. Outcome endpoints included clinical benefit rate (CBR), progression-free survival (PFS), and overall survival (OS). CBR was defined as the rate of complete response, partial response, or stable disease lasting at least six months, according to RECIST 1.1 criteria. OS and PFS were defined as the interval from treatment initiation to death and disease progression, respectively. TMB was uniformly derived from tumor sequencing data, representing the count of nonsynonymous mutations relative to the target sequencing size. TMB cutoffs were evaluated for outcome associations using study replicability analysis, alongside individual-study analysis and random-effects meta-analysis for comparison.

**RESULTS:** The data provided sufficient evidence of replicable outcome associations for specific TMB cutoffs in melanoma, lung cancer, and bladder cancer. The FDA-recommended cutoff of 10 mutations per megabase showed replicable associations in melanoma for OS (p-value < 0.01) and CBR (p-value < 0.01), though more replicable cutoffs were identified for the latter. Lower cutoffs of 4 and 2 were found to be replicable in lung cancer for CBR (p-value = 0.04) and in bladder cancer for OS (p-value < 0.01), respectively. No cutoff was deemed replicable for the other cancer type and outcome combinations, due to no association, inadequate power, or insufficient data.

**CONCLUSION:** A pan-cancer cutoff of 10 mutations per megabase may not be optimal for predicting immunotherapy outcomes. Further studies are needed to determine appropriate cutoffs specific to cancer types and outcomes through statistically principled replicability analyses.

## 1 Introduction

Tumor Mutational Burden (TMB), defined as the total number of somatic, nonsynonymous mutations in the DNA of a tumor cell, is typically measured by sequencing technologies and has emerged as a promising biomarker in cancer immunotherapy.^1–5^ It is plausible that tumors with a higher mutational burden harbor more neoantigens (that is, abnormal proteins recognized and targeted by the immune system), potentially enhancing anti-tumor immune response.^6–8^ Elevated TMB levels have been associated with improved clinical outcomes in various studies of immunotherapy.^1,2,9^ To enhance the clinical utility of TMB as an immunotherapy biomarker, efforts have been undertaken to establish a cutoff for dichotomizing TMB. Notably, the US Food and Drug Administration (FDA) approved pembrolizumab for patients with ≥10 mutations/megabase (mut/Mb).^10,11^ However, conflicting evidence persists on the cutoff choice.^12^ For example, Samstein et al (2019)^2^ suggested categorizing TMB by quantiles, while Zheng (2023)^13^ advocated for a fixed cutoff of 13 mut/Mb in select cancer types and Mo et al (2023)^14^ supported the cancer-agnostic cutoff of 10 mut/Mb.

These conflicts may have arisen from limitations such as biased representation of cancer types in the supporting data, unrealistic imposition of shared cutoffs and their effects across cancer types, and inadequate examination of data across multiple studies. More specifically, the popular cutoff of ≥10 mut/Mb, recommended by the FDA, was driven by the KEYNOTE-158 study comprising 790 patients with advanced solid tumors treated by pembrolizumab.^15^ It reported a clinical benefit rate (CBR) of 29% (95% Confidence Interval [CI]: 21% ∼ 39%) for TMB-High patients (n=102, 13%) and 6.3% (95% CI: 4.6% ∼ 8.3%) for TMB-Low patients (n=688, 87%), each including multiple cancer types. In this study, TMB-High patients were primarily diagnosed with lung, cervical, and endometrial cancer, comprising 64% of the study patients. It did not include prevalent cancer types such as prostate cancer, despite these patients still being treated based on the ≥10 cutoff.^12^ Subsequent studies sought to validate this cutoff, with some challenging its effectiveness across diverse cancer types^12,16^ and others showing its limited value as a biomarker for outcome endpoints such as progression-free survival.^17^ In a recent attempt to consolidate data across studies, Bareche et al (2022)^18^ surveyed published studies and conducted a multi-study analysis using a random-effects meta-analysis (REMA) approach,^19^ allowing for random deviations from the overall effect to accommodate study heterogeneity. However, REMA analysis may still be impacted by conflicting or outlying associations when studies are heterogeneous (as shown in Figure 1 of the Bareche et al paper^18^). Consequently, its results may mask true effects or falsely indicate associations.

**Figure 1:**
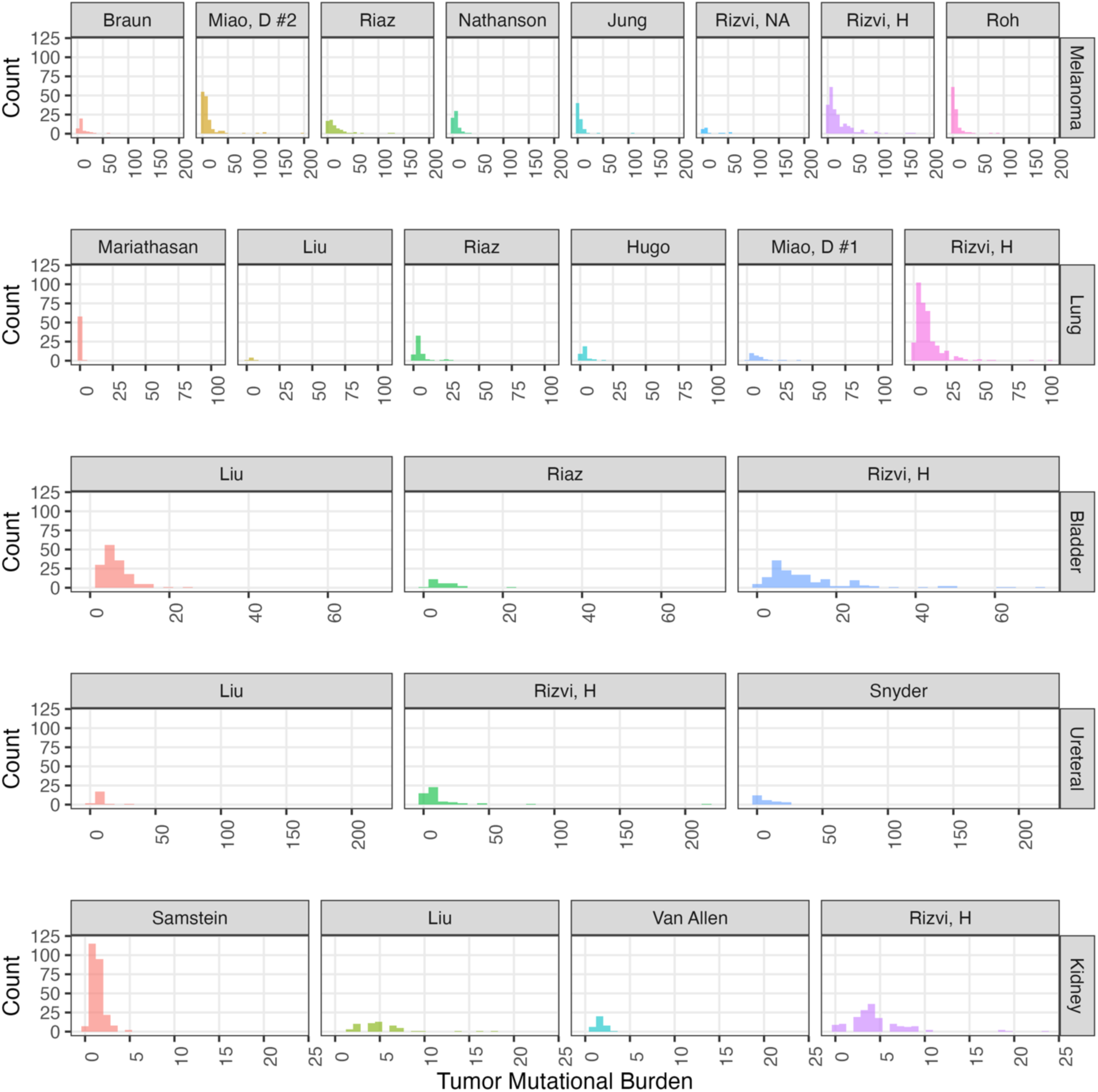
Histograms of tumor mutational burden by studies available in PredictIO for melanoma, lung, bladder, ureter, and kidney cancer.

Methodologies tailored to study replicability, such as those reviewed in Bomogolov & Heller 2023^20^ and others,^21–23^ are more suitable to explore heterogeneous effects in multi-study analyses. Unlike REMA, replicability analyses do not assume a common association across studies. They are particularly useful when outlying studies spur significance in REMA but do not ensure replicability, which is a common misconception.^18^ Over the last two decades, the replicability crisis spanning disciplines has particularly propelled the development of statistical tools to investigate study replicability.^20, 24^

We sought to utilize replicability analysis to identify TMB cutoffs with multi-study data in a cancer-specific and endpoint-aware manner. Besides relying on fewer assumptions, it is more robust to the likely heterogeneity in TMB measures across studies (which may use different sequencing platforms), as it evaluates TMB association within each study without necessitating TMB harmonization across studies. Given the relatively limited number of published TMB studies per cancer type and predominantly small sample sizes, the method introduced by Jaljuli et al (2023)^25^ is considered the most suitable for the replicability analysis, which employs an assumption-free approach to calculate a lower bound on the number of studies demonstrating clinical benefit (that is, outcome association in the beneficial direction) at a 95% CI.

## 2 Methods

### 2.1 Data

The PredictIO data was compiled by Bareche et al (2022), who conducted a literature review to identify relevant studies published between January 2015 and June 2022.^18,26^ Studies involving immune checkpoint blockade treatment in the adjuvant setting were included, excluding those that combined this treatment with chemotherapy, radiation therapy, or other molecularly targeted treatment. The included studies had to comprise at least 20 patients with tumor sequencing data (for whole exome or targeted cancer genes). TMB was derived from the sequencing data as the number of non-synonymous mutations divided by the number of the sequencing target megabases. Clinical outcomes included clinical benefit rate (CBR), progression-free survival (PFS), and overall survival (OS). CBR was defined as the rate of complete response, partial response, or stable disease lasting at least six months, according to RECIST 1.1 criteria. OS (PFS) was defined as the interval from treatment initiation to death (disease progression), with a censoring window of 36 (24) months to adjust for the variable clinical follow-up between studies. We focused on cancer types represented in at least two studies for a given outcome.

### 2.2 Single-Cohort Analysis

Each potential TMB cutoff, ranging from 1 to 20, was tested for association with an available outcome in each cohort (defined as each combination of a published study and a cancer type) within PredictIO. Throughout this article, we refer to the single-study analysis for each ‘cohort’ as the single-cohort analysis. At each cutoff *c*, patients were categorized as TMB-High if ***TMB*** ≥ *c*, and TMB-Low otherwise. Outcomes were compared between TMB-High and TMB-Low groups in each cohort, with the odds ratio (OR) estimated for CBR and hazard ratios (HRs) for OS and PFS. The analyses were performed using R version 4.3.2, R package ‘stats’ (version 4.3.2) for logistic regression, and R package ‘survival’ (version 3.5-7) for survival regression.^27,28^

### 2.3 Random-Effects Meta-Analysis

Meta-analysis models are widely adopted for aggregating results of independent studies in systematic reviews and are particularly useful for drawing conclusion from small studies with weak signals.^29,30^ Random-Effects Meta-Analysis (REMA) is frequently employed to accommodate signal heterogeneity, by assuming that the study-specific effects come from a Gaussian distribution (see notations and equations in Text Box 1).^31,32^ We conducted REMA with the PredictIO data both by individual cancer types and for all available cancer types combined, using the R package ‘meta’ (version 6.5-0).^33^ The former analysis is for comparison with other cancer-specific analysis conducted in our study and the latter is to bridge with the pan-cancer REMA reported in the literature.^18^

### 2.4 Replicability Analysis

We applied the method by Jaljuli et al (2023)^25^ to assess the replicability of outcome association findings across studies for each cancer type, avoiding the assumption of a common effect shared among them. Some cutoffs may be extreme for cohorts with limited sample sizes, creating groups with too few patients or events. Such cohorts were excluded from replicability analysis, as this analysis required individual-study summary statistics. In Text Box 1, we motivated the construction of the suggested replicability metric, denoted by **u_adj_(*c*)**, which is the estimated 95% confidence lower bound on the number of studies replicating an association effect adjusted for the total number of analyzed cohorts, while penalizing on the number of cohorts lost when dichotomizing.^25^ This metric ranges from 0 to 1, where 1 indicates complete replicability and 0 signifies a failure to meet the minimal requirement for replicability (that is, only two studies have an association effect replicated). For cancer types with multiple replicable cutoffs nominated, the cutoff that maximized the metric was preferred. Replicability analysis was performed using the R package ‘metarep’ (version 1.2) with the default truncation parameter *t* = 0.2 and a one-sided 95% confidence level.

**Text Box 1:**
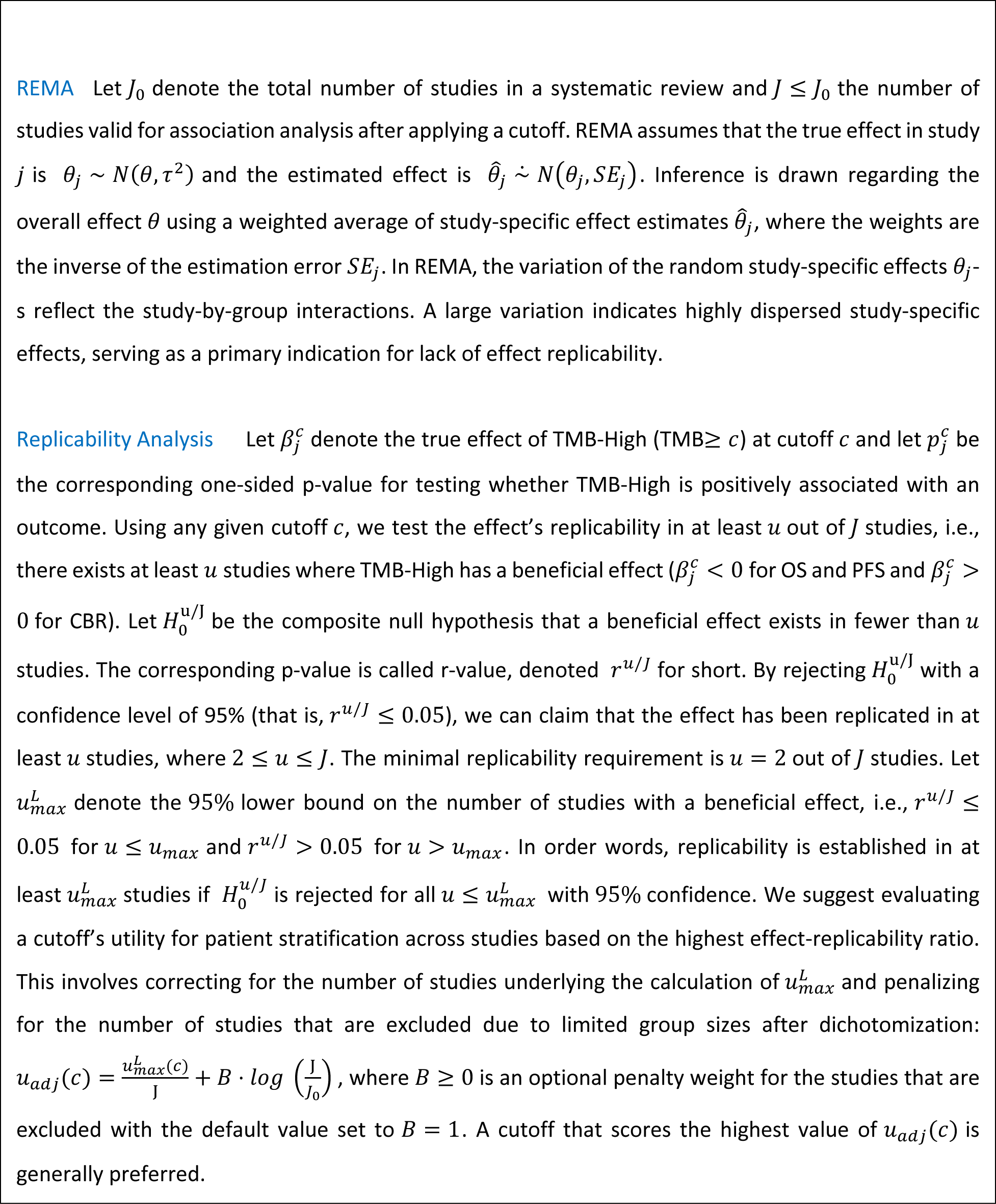
Notations and Equations for REMA and Replicability Analysis.

## 3 Results

### 3.0 TMB Distribution by Cancer Type

The TMB level ranged widely from 0.01 to 196 mut/Mb and varied by cancer type (Figure 1 and Figure S1). It skewed towards smaller values in kidney cancer (median [IQR]: 2 [1-4]), compared to melanoma (6 [2-14]), lung (6 [4-11]), bladder (6 [4-10]), and ureter cancer (5 [3,9]) (Table S1). Consequently, what constituted a high TMB level in one cancer type may be considerably lower than others. Among the 495 patients with kidney cancer, only 2% qualified as TMB-High at the 10 mut/Mb cutoff. In contrast, among 727 patients with melanoma, 34% surpassed this cutoff.

### 3.1 Single-Cohort Analyses

Single-cohort analyses demonstrate a strong dependence of statistical association on not only the TMB cutoffs, but also the cancer types and outcome endpoints (Figure 2).

**Figure 2:**
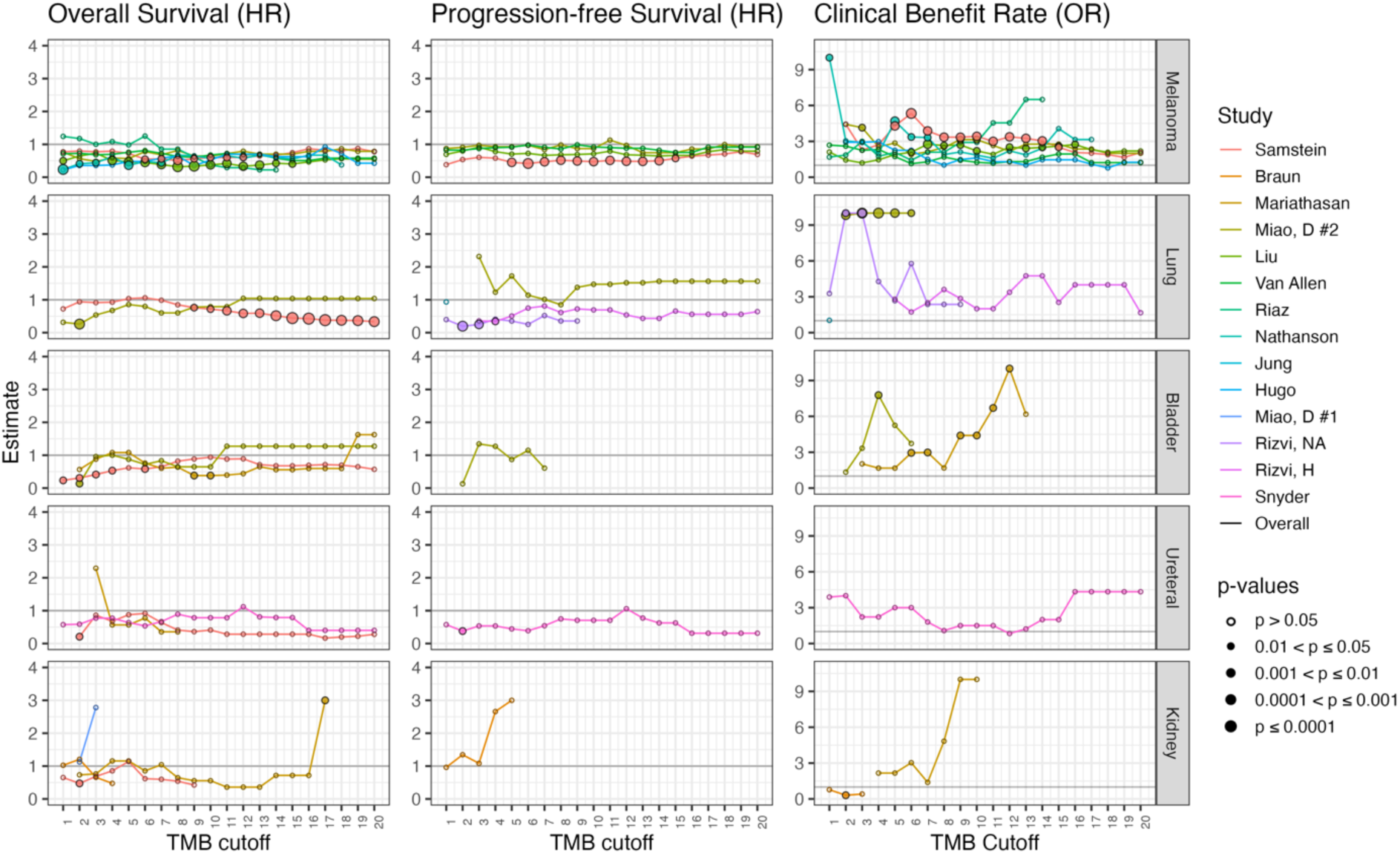
Estimated effects for the single-cohort analysis on the association of TMB-High (***TMB*** ≥ ***c***) with overall survival (left column of panels), progression-free survival (middle column), and clinical benefit rate (right column), along with their corresponding p-values (indicated by point size and type), as a function of the cutoff ***c***. For each outcome endpoint, analyses were performed for each study (indicated by colors) of five individual cancer types (as five panel rows) included in at least two studies within PredictIO.

Among the melanoma cohorts, statistical significance was observed in 3 cohorts for OS, 1 cohort for PFS, and 4 cohorts for CBR at one or more TMB cutoffs. In the lung cancer cohorts, significance was found in 2 cohorts each for OS, PFS, and CBR. For bladder cancer, significance was seen in 3 cohorts for OS and 2 cohorts for CBR, while for kidney cancer, significance was noted in 2 cohorts for OS and 1 cohort for CBR. No significance was found for PFS in either bladder or kidney cancer cohorts. In the other cancer types, for all three endpoints, significance was found in only one cohort or not at all at any TMB cutoff.

For all cancer types and endpoints, the estimated ORs for CBR and HRs for OS and PFS exhibited non-monotonic behavior as the cutoff increased, showing substantial fluctuations in effect size and occasional changes in effect sign. Of note, at the 10 mut/Mb cutoff, statistical significance was seen only in a minority of cohorts, including 4 out of 15, 1 out of 7, and 4 out of 11 cohorts for OS, PFS, and CBR, respectively.

### 3.2 Random-Effects Meta-Analyses

In a REMA that aggregate the data across the available studies (regardless of cancer types), significant effects were found at all examined cutoffs for OS, at all but one cutoff for CBR (*c* ≥ 1), and at most cutoffs for PFS (3 ≤ *c* ≤ 16) (Figure 3.A). This result underscores the lack of a statistical basis for favoring the FDA-recommended TMB cutoff of 10 mut/Mb over other values, which exhibited comparable statistical significance. This result also highlights the problematic sensitivity of REMA to a single outlying study, as for example PFS association was reported in no but one (large) individual study for cutoffs from 5 to 15 (Figure S2).

**Figure 3.**
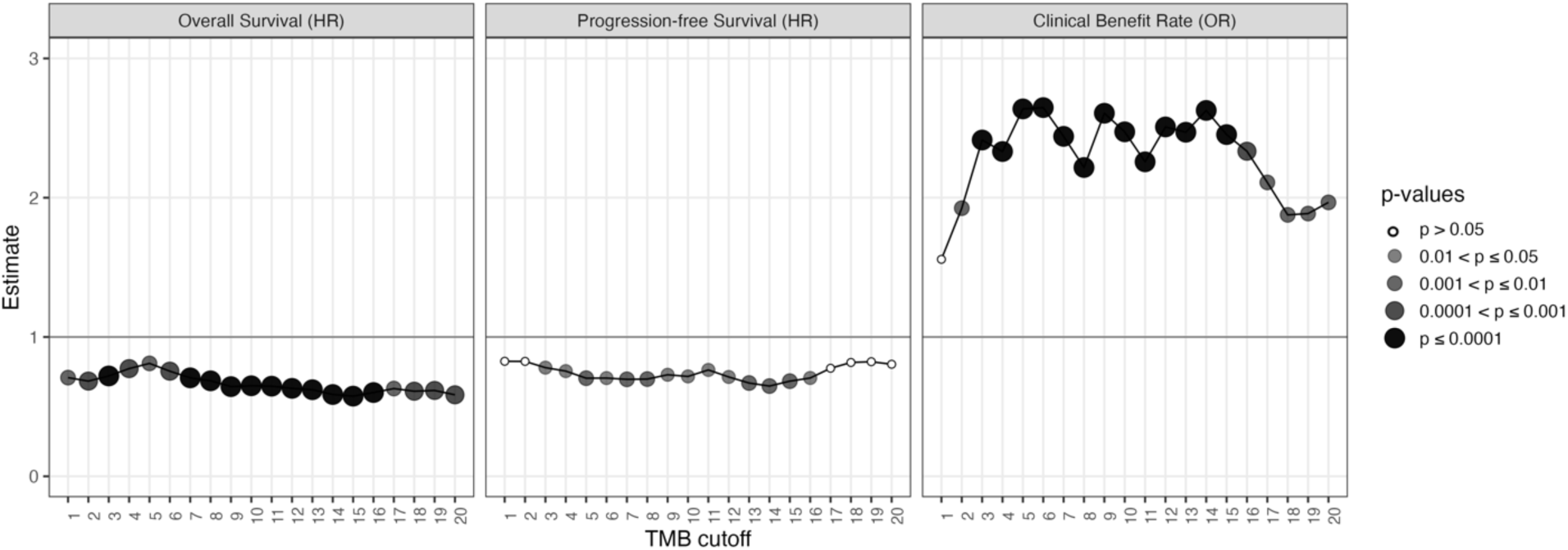
**A:** Estimated effects for the random-effects meta-analysis on the association of TMB-High (***TMB*** ≥ ***c***) with overall survival (left column of panels), progression-free survival (middle column), and clinical benefit rate (right column), along with their corresponding p-values (indicated by point size and type), as a function of the cutoff ***c***. For each outcome endpoint, analyses were performed across all available studies within PredictIO regardless of cancer types.

Figure 3.B shows the results of REMAs aggregating the data across cohorts in a cancer-specific manner.

**Figure 3.**
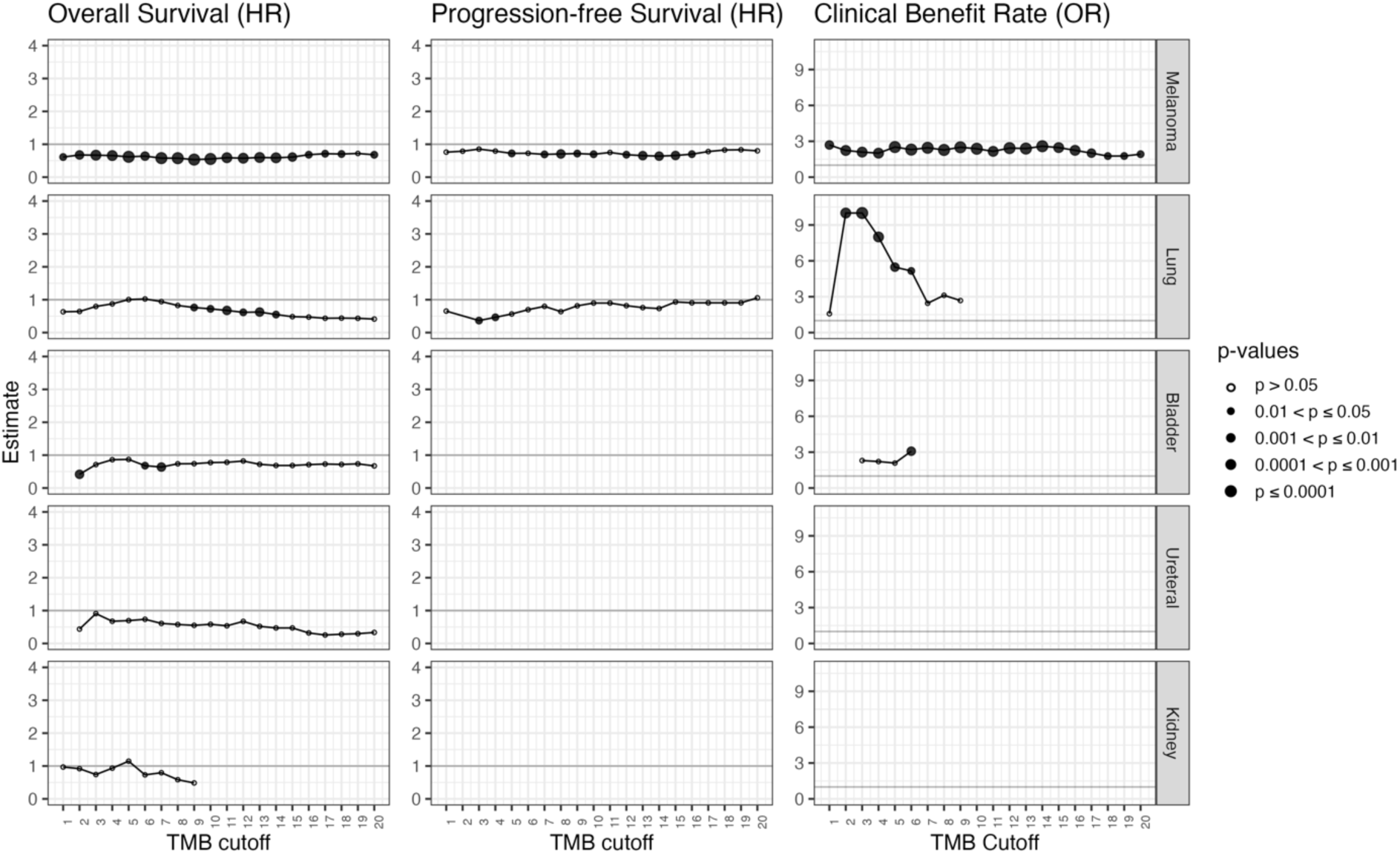
**B:** Estimated effects for the random-effects meta-analysis on the association of TMB-High (***TMB*** ≥ ***c***) with overall survival (left column of panels), progression-free survival (middle column), and clinical benefit rate (right column), along with their corresponding p-values (indicated by point size and type), as a function of the cutoff ***c***. For each outcome endpoint, analyses were performed for the five individual cancer types (as five panel rows) included in at least two studies within PredictIO.

TMB-High at the 10 mut/Mb cutoff exhibited significant outcome associations only in melanoma (with all three endpoints) and lung cancer (with OS).

Broader cancer-specific REMAs across all possible TMB cutoffs showed a significant association with OS in melanoma at all cutoffs except for one, despite limited significance in single-cohort analyses across all seven available cohorts. REMAs of OS in lung cancer reported significant p-values at six cutoffs, with significance predominantly driven by the considerably larger cohort among the two analyzed (355 patients in Samstein et al 2019^2^ *versus* 57 patients in Miao et al 2018^34^). Additionally, significance was seen for OS REMAs in bladder cancer at a small number of low-valued cutoffs. No cutoff was deemed to be associated with a significant effect in kidney and ureter cancer for OS.

Similar broad REMAs for PFS showed significance at half of the examined cutoffs in melanoma, primarily driven by the largest cohort among the four available (as indicated by the insignificant single-cohort analyses in the remaining three cohorts and the insignificant REMA when excluding the largest cohort^2^). PFS REMAs also identified statistical significance in lung cancer at only two small cutoffs (2 and 3 mut/Mb). Data was insufficient to conduct PFS REMAs in bladder, kidney, and ureter cancer.

For CBR, broad cancer-specific REMAs found statistical significance again in melanoma across all examined cutoffs, and sporadic significance at lower cutoffs in lung and bladder cancer. Data was insufficient for such analyses in kidney cancer and ureter cancer.

### 3.3 Replicability Analyses

For each cancer type, we assessed replicability using the ***u***_adj_ (*c*) metric at all possible cutoffs (Figure 4 and Table S2).

**Figure 4:**
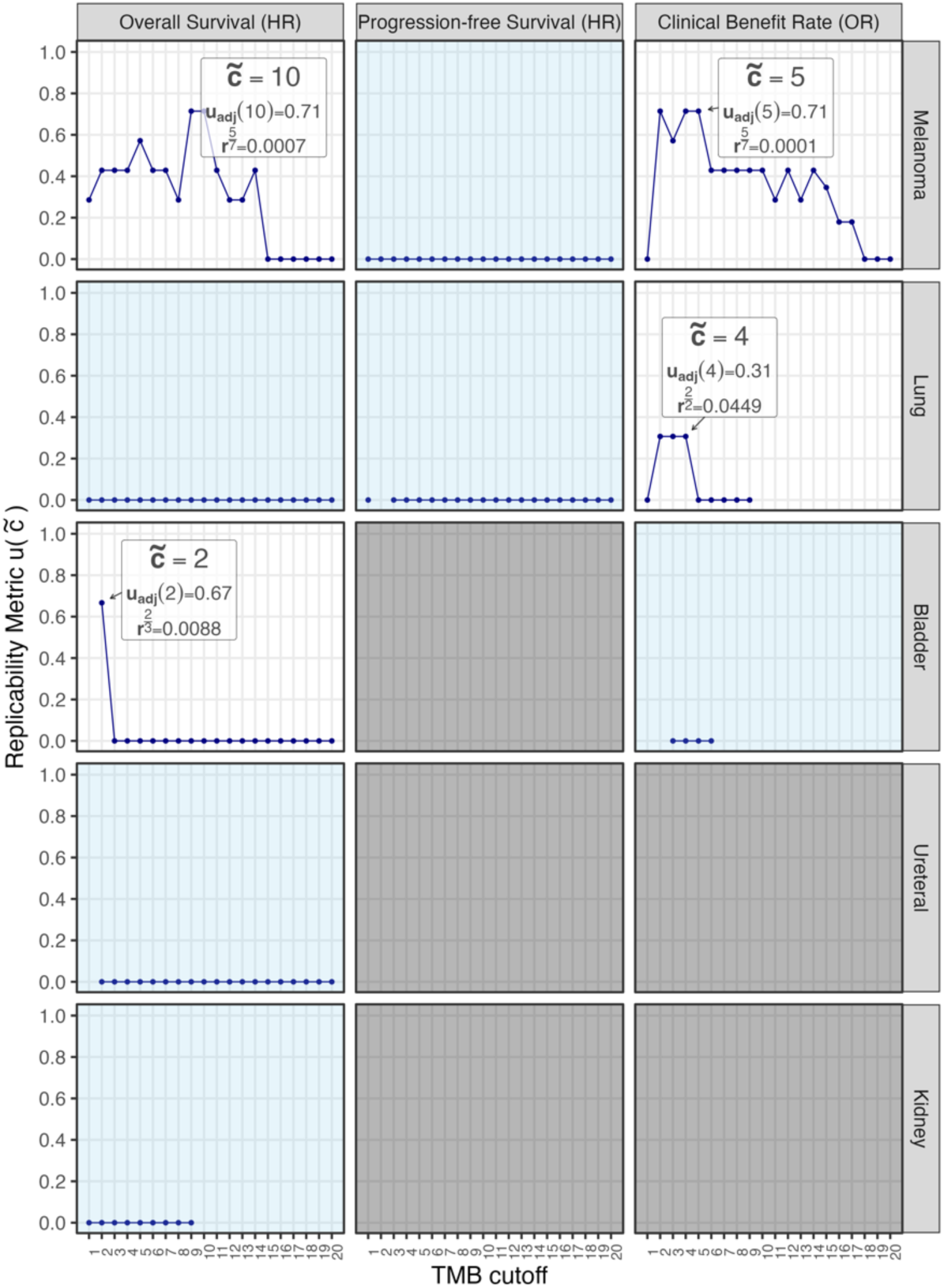
Estimated metric ***u***_***adj***_(***c***) for the replicability analysis on the association of TMB-High (***TMB*** ≥ ***c***) with overall survival (left column of panels), progression-free survival (middle column), and clinical benefit rate (right column), as a function of the cutoff ***c***, in each of five individual cancer types (as five panel rows). For each outcome endpoint in each cancer type, a cutoff was (1) either nominated based on the highest replicability metric, (2) not nominated due to no evidence of replicability (that is, ***u*<2** across the studies analyzed) at all cutoffs (indicated by blue panels), or (3) not nominated due to insufficient data (that is the number of studies with estimable effects is less than two) for all cutoffs (indicated by grey panels). When nominated, the cutoff ***c***V was selected based on the highest ***u***_***adj***_(***c***) metric value. Both values, along with the corresponding r-value, are indicated in the panel.

This analysis supported that TMB-High, defined by the 10 mut/Mb cutoff, was associated with OS in melanoma, with a significant replicability rate (p-value of *r*^5/7^ < 0.001). The cutoff of 5 achieved the most replicable effect for CBR in melanoma, with a significant replicability rate (p-value of *r*^5/7^ < 0.001). However, no replicable cutoff can be found for PFS in melanoma.

In lung cancer, the cutoff of 4 was found to be significantly replicable for CBR (p-value of *r*^2/2^ = 0.045). Comparable replicability rates were reported at *c* = 2 and *c* = 3 with lower p-values (0.009 and 0.003, respectively). However, *c* = 4 was preferred for its better balance in patient numbers, potentially offering more clinical utility. No replicable cutoffs were established for OS and PFS in lung cancer.

In bladder cancer, *c* = 2 was nominated for OS, demonstrating replicated association in 2 out of 3 cohorts with statistical significance (p-value of *r*^2/3^ = 0.009). Replicability assessment was not possible for PFS in bladder cancer due to data insufficiency, where in CBR, effect replicability was not established across any cutoff.

In both ureteral cancer and kidney cancer, no replicable cutoffs were established for OS, and no sufficient data was available to assess replicability for PFS and CBR.

## 4 Discussion

The literature suggests an association between higher TMB levels and better immunotherapy outcomes, followed by the dichotomization of TMB levels for enhanced clinical utility.^35^ Despite information loss associated with dichotomization, this approach has gained traction for its practicality in a clinical setting. Diverse cutoffs for TMB dichotomization have been suggested in a cancer-type-agnostic manner without robust statistical evidence. To address this important and pressing issue, we performed replicability analysis for multi-study data in a cancer-specific and outcome-aware manner.

Our cancer-specific analysis revealed substantial variation in TMB distributions, association effects, and selected cutoffs across cancer types, affirming the need for a cancer-type-specific approach. Our results raised concerns about the validity of the FDA-recommended TMB-High cutoff of 10 mut/Mb as a *de facto* rule across cancer types. Firstly, TMB varies significantly across cancer types, ranging from as low as 0.01 mut/Mb to nearly 200 mut/Mb in the data we studied. Within this wide range, an arbitrary selection of TMB cutoffs can restrict its clinical utility. Secondly, statistical significance of the combined effect using REMA was observed in only two cancer types, suggesting a potential lack of association or an insufficiency of statistical power in the other cancer types, and leaving the magnitude of this association unknown for many other cancer types. Thirdly, replicability could not be established for less studied cancer types (that is, bladder and ureteral cancer) and for the more clinically relevant endpoints in certain types (that is, PFS and OS for lung cancer), for which we recommend additional well-designed studies to address data limitations and enable cutoff nomination.

Our analysis underscored the importance of identifying cutoffs with the highest replicability, echoing the FDA requirement for replicability that mandates at least two studies reporting significant p-values.^36^ Despite the acknowledged limitations of this requirement,^25^ it stresses the imperative for additional validation studies, which was precisely corroborated by our results. In this study, we implemented an assumption-free version of replicability analysis based on directional p-values (or equivocally, signed effect sizes, as p-values are a one-to-one transformation of effect sizes). By utilizing p-values for further analyses, we enable robust inference without imposing additional assumptions, thereby gaining robustness and power compared to parametric methods. We acknowledge the risks associated with the use of p-values, as highlighted by recent publications, including a statement by the ASA board^37^. However, we emphasize that these risks stem from the misuse of the p-values rather than flaws inherent to the measure itself^38^. Directly connected to p-values, estimated effect sizes and confidence intervals are subject to the same potential misuse when tied to statistical significance. As Benjamini (2016)^38^ noted, “*In some sense it* [the p-value*] offers a first line of defense against being fooled by randomness, separating signal from noise, because the models it requires are simpler than any other statistical tool needs*.”

Our study also highlighted the limitations of relying solely on statistical significance from REMA, which is a commonly used and potentially powerful approach. Two key limitations of REMA are: (1) cross-study analysis may be driven by a single outlier study, as there is no definitive rule in practice for identifying and excluding outlier studies, and (2) statistical significance at a specific cutoff does not necessarily indicate that it is the best cutoff. Furthermore, the replicability approach provided greater power than the REMA approach. For example, our study revealed a replicable effect for OS in bladder cancer at the cutoff of 2 using the former approach but not the latter.

In this article, we delineated between the term ‘replicability’ from ‘reproducibility,’ which are often used interchangeably without clear distinction. Following editorials in the Biostatistics journal, ^39,40^ we made a crucial distinction: reproducibility pertains to the property of a study, signifying the possibility for others to recreate the specific analyses and findings using the original data and methods; in contrast, replicability refers to the ability to obtain consistent and similar results when an entire study — including enlisting subjects, collecting data, and producing analysis results — is conducted anew in a similar but not necessarily identical manner, yet achieving comparable results. ^41,42^ While reproducibility is important for validating the methods and analyses within a study, replicability extends this concept to the broader context of being able to validate the findings in additional studies, often using different samples or settings and thus not dependent on specific contextual factors. One should ensure study reproducibility and strive to improve study replicability.

Upon the suggestion of a reviewer, we would also like to comment on replicability (narrow defined studies) versus generalizability (broadly defined studies). In practice, these definitions largely depend on the granularity of study parameters, such as patient population, treatment regime, reference control, and outcome. This level of granularity has not been adequately addressed in the still young and evolving field of cancer immunotherapy and biomarker development. Unlike many published TMB studies, which often assume the universal applicability of a single cutoff across diverse contexts, a key finding of our work is that optimal TMB cutoffs may vary depending on the tumor type and the outcome being studied. In other words, the primary objective of our manuscript is to assess the replicability of dichotomized TMB as a biomarker within specific clinical contexts, defined by combinations of tumor type and outcome. While our study does not further refine analyses based on treatment regime and control selection due to limited number of available studies, our findings lay the groundwork for advancing the needed granularity in cancer immunotherapy research and development.

Our analysis does not distinguish whether TMB serves as a prognostic or predictive biomarker, nor does it account for other potential moderators of cancer immunotherapy, such as PD-L1 expression and interferon gene signatures, primarily due to data limitations. Research interest in TMB was initially sparked within the context of immunotherapy, based on its plausible underlying biological mechanism. The potential for TMB to act as a prognostic biomarker is likely context-dependent and should be explored further in carefully designed studies involving non-immunotherapy treatments.

In conclusion, our study highlights the importance of adopting a nuanced, cancer-specific approach to determining TMB cutoffs by outcome endpoint, guided by replicable evidence from multi-study data. Further validation studies, including randomized controlled studies that are carefully designed using empirical evidence from observation studies and non-randomized trials, across diverse cancer types are warranted to enhance clinical utility and ensure replicable research findings in the field of cancer immunotherapy.

## Author Contributions

Concept and design: Qin, Jaljuli Acquisition, analysis, or interpretation of data: Qin, Jaljuli, Whiting Drafting of the manuscript: Qin, Jaljuli Critical review of the manuscript for important intellectual content: All authors Statistical analysis: Jaljuli, Whiting Obtained funding: Qin.

## Data and Code Availability

The R code used for our analyses can be freely downloaded from GitHub (https://github.com/LXQin/TMB_Cutpoint_Nomination). The PredictIO data is publicly available on the Zenodo platform (https://doi.org/10.5281/zenodo.6142357) as described in its original publication.

---

**Figure S1:**
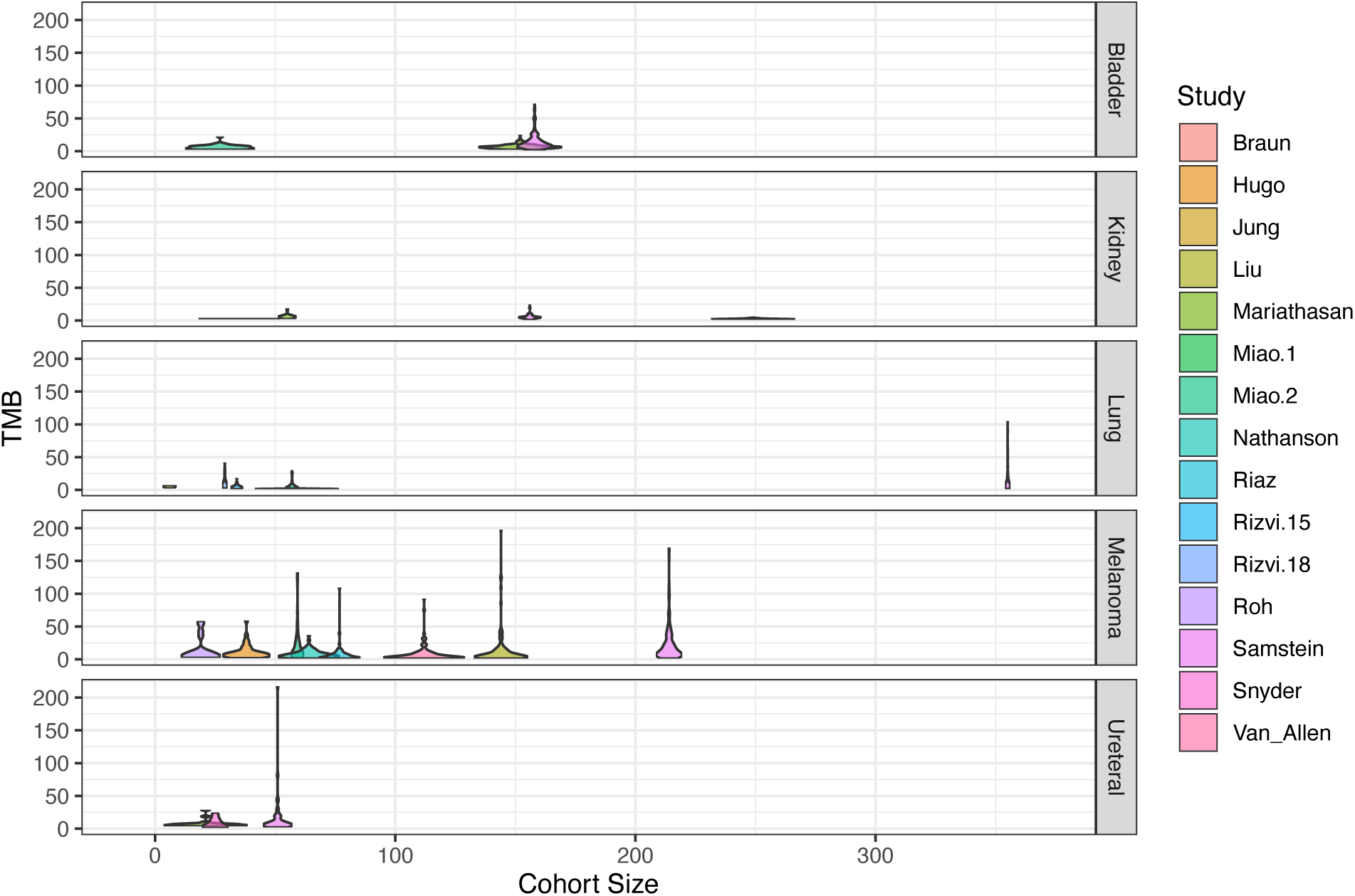
Boxplots of tumor mutational burden (original scale) by studies available in PredictIO for melanoma, lung, bladder, ureter, and kidney cancer. The studies are sorted by their respective sample sizes.

**Figure S2:**
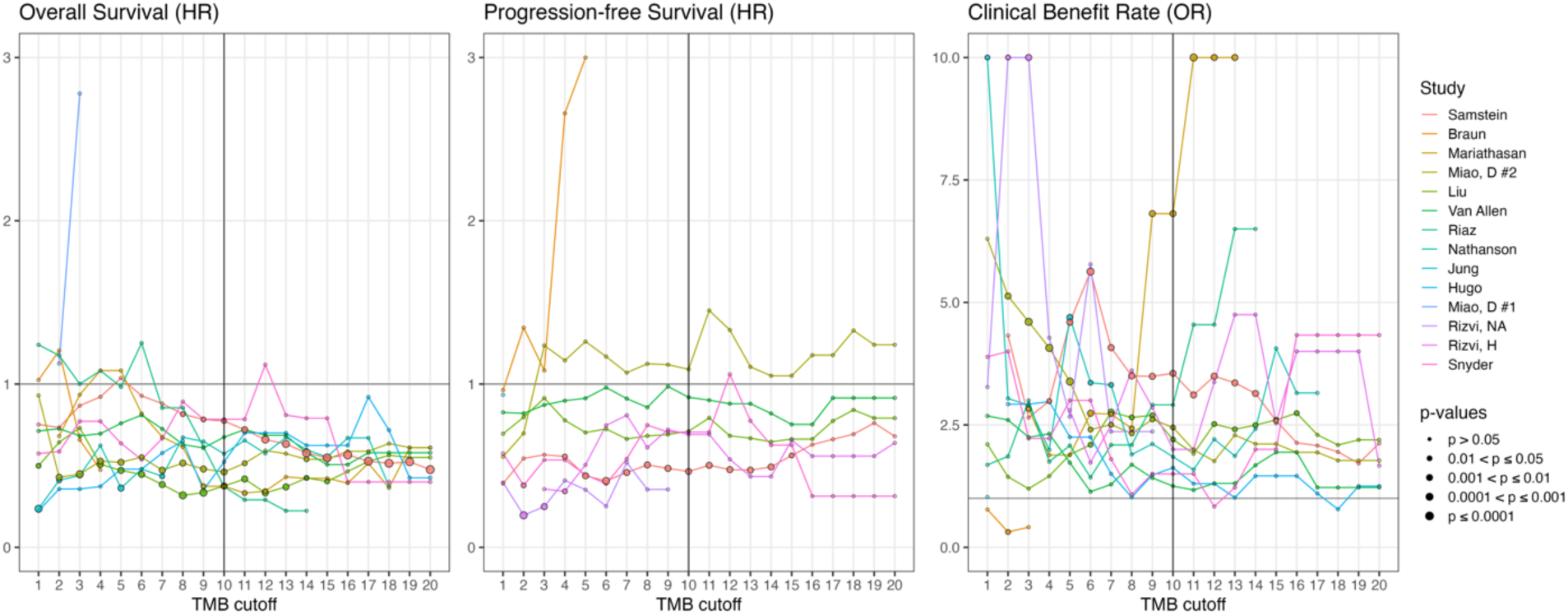
Estimated effects for the single-study analysis (regardless of cancer type) on the association of TMB-High (***TMB*** ≥ ***c***) with overall survival (left column of panels), progression-free survival (middle column), and clinical benefit rate (right column), along with their corresponding p-values (indicated by point size and type), as a function of the cutoff ***c***. For each outcome endpoint, analyses were performed for each study (indicated by colors), regardless of cancer type, included in at least two studies within PredictIO.

## References

1. Goodman AM, Kato S, Bazhenova L, et al. Tumor mutational burden as an independent predictor of response to immunotherapy in diverse cancers. Molec Cancer Therap. 2017;16(11):2598–2608.

2. Samstein RM, Lee CH, Shoushtari AN, et al. Tumor mutational load predicts survival after immunotherapy across multiple cancer types. Nat Genet. Feb 2019;51(2):202–206. doi:10.1038/s41588-018-0312-8

3. Chan, TA. TMB as a Biomarker for Immunotherapy. Cancer Cell. 2019; 201936(6), 637–638. DOI: 10.1016/j.ccell.2019.11.002.

4. Ricciuti B, Wang X, Alessi JV, et al. Association of High Tumor Mutation Burden in Non-Small Cell Lung Cancers With Increased Immune Infiltration and Improved Clinical Outcomes of PD-L1 Blockade Across PD-L1 Expression Levels. JAMA Oncol. Aug 1 2022;8(8):1160–1168. doi:10.1001/jamaoncol.2022.1981

5. Graf RP, Fisher V, Huang RSP, et al. Tumor Mutational Burden as a Predictor of First-Line Immune Checkpoint Inhibitor Versus Carboplatin Benefit in Cisplatin-Unfit Patients With Urothelial Carcinoma. JCO Precis Oncol. 2022;(6):e2200121. doi:10.1200/po.22.00121

6. Snyder A, Makarov V, Merghoub T, et al. Genetic basis for clinical response to CTLA-4 blockade in melanoma. N Engl J Med. Dec 4 2014;371(23):2189–2199. doi:10.1056/NEJMoa1406498

7. Schumacher TN, Schreiber RD. Neoantigens in cancer immunotherapy. Science. 2015;348(6230):69–74.

8. McGranahan N, Furness AJ, Rosenthal R, et al. Clonal neoantigens elicit T cell immunoreactivity and sensitivity to immune checkpoint blockade. Science. Mar 25 2016;351(6280):1463–9. doi:10.1126/science.aaf1490

9. Iams WT, Porter J, Horn L. Immunotherapeutic approaches for small-cell lung cancer. Nat Rev Clin Oncol. 2020;17(5):300–312.

10. Marcus L, Fashoyin-Aje LA, Donoghue M, et al. FDA approval summary: pembrolizumab for the treatment of tumor mutational burden–high solid tumors. Clin Cancer Res. 2021;27(17):4685–4689.

11. US Food and Drug Administration. (2020). FDA Approves Pembrolizumab for First-Line Treatment of Head and Neck Squamous Cell Carcinoma. [Press release]. Retrieved from https://www.fda.gov/drugs/drug-approvals-and-databases/fda-approves-pembrolizumab-first-line-treatment-head-and-neck-squamous-cell-carcinoma.

12. Prasad V, Addeo A. The FDA approval of pembrolizumab for patients with TMB >10 mut/Mb: was it a wise decision? No. Ann Oncol. Sep 2020;31(9):1112-1114. doi:10.1016/j.annonc.2020.07.001

13. Zheng M. Tumor mutation burden for predicting immune checkpoint blockade response: the more, the better. J Immunother Cancer. Jan 2022;10(1)doi:10.1136/jitc-2021-003087

14. Mo SF, Cai ZZ, Kuai WH, Li X, Chen YT. Universal cutoff for tumor mutational burden in predicting the efficacy of anti-PD-(L)1 therapy for advanced cancers. Front Cell Dev Biol. 2023;11:1209243. doi:10.3389/fcell.2023.1209243

15. Marcus L, Fashoyin-Aje LA, Donoghue M, et al. FDA Approval Summary: Pembrolizumab for the Treatment of Tumor Mutational Burden-High Solid Tumors. Clin Cancer Res. Sep 1 2021;27(17):4685–4689. doi:10.1158/1078-0432.CCR-21-0327

16. McGrail D, Pilié P, Rashid N, et al. High tumor mutation burden fails to predict immune checkpoint blockade response across all cancer types. Annals of Onc. 2021;32(5):661–672.

17. Aggarwal C, Ben-Shachar R, Gao Y, et al. Assessment of Tumor Mutational Burden and Outcomes in Patients With Diverse Advanced Cancers Treated With Immunotherapy. JAMA Netw Open. May 1 2023;6(5):e2311181. doi:10.1001/jamanetworkopen.2023.11181

18. Bareche Y, Kelly D, Abbas-Aghababazadeh F, et al. Leveraging big data of immune checkpoint blockade response identifies novel potential targets. Ann Oncol. Dec 2022;33(12):1304–1317. doi:10.1016/j.annonc.2022.08.084

19. DerSimonian R, Laird N. Meta-analysis in clinical trials. Control Clin Trials. Sep 1986;7(3):177–88. doi:10.1016/0197-2456(86)90046-2

20. Bogomolov M, Heller R. Replicability across multiple studies. Stat Sci. 2023;38(4):602–620.

21. Zhang J, Fu H, Carlin BP. Detecting outlying trials in network meta-analysis. Stat Med. Aug 30 2015;34(19):2695–707. doi:10.1002/sim.6509

22. Makinde FL, Tchamga MSS, Jafali J, Fatumo S, Chimusa ER, Mulder N, Mazandu GK. Reviewing and assessing existing meta-analysis models and tools. Brief Bioinfo. 2021;22(6)doi:10.1093/bib/bbab324

23. Xiao M, Chu H, Hodges JS, Lin L. Quantifying replicability of multiple studies in a meta-analysis. Annals of App Stat. 2024;18(1):664-682, 19.

24. PSYCHOLOGY. Estimating the reproducibility of psychological science. Science. Aug 28 2015;349(6251):aac4716. doi:10.1126/science.aac4716

25. Jaljuli, I., Benjamini, Y., Shenhav, L., Panagiotou, O. A., & Heller, R. (2023). Quantifying Replicability and Consistency in Systematic Reviews. Stats in Biopharm Res. 15(2), 372–385. 10.1080/19466315.2022.2050291.

26. Bareche, Y. (2022). Zenodo. barechey/PredictIO.data: (v2.0) [Data set]. 10.5281/zenodo.6142357.

27. Terry M. Therneau, Patricia M. Grambsch (2000). _Modeling Survival Data: Extending the Cox Model_. Springer, New York. ISBN 0–387-98784-3.

28. Therneau T (2024). A Package for Survival Analysis in R. R package version 3.5–8, https://CRAN.R-project.org/package=survival.

29. Burke DL, Ensor J, Riley RD. Meta-analysis using individual participant data: one-stage and two-stage approaches, and why they may differ. Stat Med. Feb 28 2017;36(5):855–875. doi:10.1002/sim.7141

30. Deeks JJ, Higgins JP, Altman DG, Cochrane Statistical Methods Group. Analysing data and undertaking meta-analyses. Cochrane handbook for systematic reviews of interventions. 2019 Sep 23:241–84.

31. Kafkafi N, Golani I, Jaljuli I, et al. Addressing reproducibility in single-laboratory phenotyping experiments. Nat Met. 2017;14(5):462–464.

32. Jaljuli I, Kafkafi N, Giladi E, et al. A multi-lab experimental assessment reveals that replicability can be improved by using empirical estimates of genotype-by-lab interaction. PLoS biology. 2023;21(5):e3002082.

33. Balduzzi S, Rücker G, Schwarzer G. How to perform a meta-analysis with R: a practical tutorial. BMJ Ment Health. 2019;22(4):153–160.

34. Miao D, Margolis CA, Vokes NI, et al. Genomic correlates of response to immune checkpoint blockade in microsatellite-stable solid tumors. Nat Genet. Sep 2018;50(9):1271–1281. doi:10.1038/s41588-018-0200-2

35. Chan TA, Yarchoan M, Jaffee E, Swanton C, Quezada SA, Stenzinger A, Peters S. Development of tumor mutation burden as an immunotherapy biomarker: utility for the oncology clinic. Ann Oncol. Jan 1 2019;30(1):44–56. doi:10.1093/annonc/mdy495

36. MDI, Metered Dose Inhaler and Drug, Dry Powder Inhaler DPI. (1998). Guidance For Industry. Center for Drug Evaluation and Research (CDER), 1000.

37. Wasserstein RL, Lazar NA. The ASA Statement on P-values: Context, Process, and Purpose. The American Statistician. 2016;70(2):129–133. doi:10.1080/00031305.2016.1154108

38. Benjamini, Y. It’s Not the P-values’ Fault. The American Statistician. 2016;70: Online Supplement to ASA Statement on P-values. available at https://www.tandfonline.com/action/downloadSupplement?doi=10.1080%2F00031305.2016.1154108&file=utas_a_1154108_sm5354.pdf.

39. Diggle PJ, Zeger SL. Editorial. Biostatistics. 2010;11(3):375-375. doi:10.1093/biostatistics/kxq029

40. Peng RD. Reproducible research and Biostatistics. Biostatistics. 2009;10(3):405-408. doi:10.1093/biostatistics/kxp014

41. Announcement: Reducing our irreproducibility. Nature. 2013/04/01 2013;496(7446):398-398. doi:10.1038/496398a

42. National Academies of Sciences E, Medicine. Reproducibility and Replicability in Science. The National Academies Press; 2019:256.

